## Supplementary Figures and Legend for "Spatial transcriptomics in the adult *Drosophila* brain and body"

### Supplementary Figure Legends

**Figure 1 – figure supplement 1: Expression of the here selected marker genes in single-cell datasets**.
(**A**) Heatmap showing expression in standardized log(CPM) of the 100 marker genes used for in the head dataset in the single-cell clusters. (**B**) Density plot showing levels of gene expression detected in scRNA-seq (Pech et al., 2024). Blue: all genes, red: genes used in the spatial study. (**C**) Heatmap showing expression in standardized log(CPM) of the 50 marker genes used for body dataset in the Fly Cell Atlas (FCA) single-cell clusters. (**D**) Density plot showing levels of gene expression detected in scRNA-seq (FCA, (Li et al., 2022)). Blue: all genes, red: genes used in the spatial study.

**Figure 1 – figure supplement 2: Head samples.**
(**A**) All detected mRNA molecules color coded in all 13 head samples used in the analysis. Scale bars represent 100 µm*.* Background stain labels DAPI. (**B**) Boxplot showing gene expression (number of detected mRNA molecules) in each of the head samples. Boxplot shows the median (center line), upper and lower quartiles (box limits) and 1.5× interquartile range (whiskers). All data points are shown. (**C**) Heatmap showing Pearson correlation between the individual head section samples and to the scRNA data from the FCA.

**Figure 1 – figure supplement 3: Body samples.**
(**A**) All detected mRNA molecules color coded for all body samples used in the analysis. Sections are from the same male. Anterior is to the left, dorsal is up on each section and the section are ordered (top to bottom). Scale bars represent 100 µm. Background stain labels DAPI. (**B**) Boxplot showing gene expression (number of detected mRNA molecules) in each of the body samples. Boxplot shows the median (center line), upper and lower quartiles (box limits) and 1.5× interquartile range (whiskers). All data points are shown. (**C**) Heatmap showing Pearson correlation between the individual samples and to the scRNA data from the FCA.

**Figure 2 – figure supplement 1: Motor neuron markers in flight muscles.**
(**A**, **B**) Two different thorax section are shown on the left and on the right. Molecular Cartography of neuronal markers in flight muscles (A). Molecular Cartography of neurotransmitter genes in flight muscles (B). (*VAChT*: cholinergic, *VGlut*: glutamatergic, *Gad1*: GABAergic). Scale bars represent 100 µm.

**Figure 2 – figure supplement 2: Comparison of body spatial datasets with body single-cell datasets.**
(**A**) Composite heatmap showing gene-gene co-expression based on Pearson correlation. Bottom triangle calculated on spatial datasets (using grid-based 5 µm squares). Top triangle calculated using single-cell data. (**B**) Gene-gene correlation measured across grid-based 5 µm squares and cells.

**Figure 2 – figure supplement 3: Colocalization of gene expression in the body datasets.**
(**A**) Overview of colocalization algorithm. For each mRNA spot a disk of 4 µm diameter was used as search space. Overlaps in the disk area were then used to calculate proximity between genes as defined by the formula in the figure. (**B**) Heatmap showing co-occurrence of genes with each other. (**C**) Ward's clustering of gene-gene proximities. (left) Dendrogram

clustering of genes. (right) Spatial location of gene clusters. Colors in the dendrogram are used to show the expression of the respective genes on the fly section.

**Figure 3 – figure supplement 1: Subcellular mRNA localization in leg and head muscles.** (**A**) Molecular Cartography of *sls, Act79B, Tpn41C* and *Mhc* mRNAs in leg muscles, showing nuclei enrichment for *sls* mRNA and proximity of *Act79B* mRNA to leg muscle nuclei. (**B**) Molecular Cartography of *sls* and *Mhc* mRNAs in head muscles, showing nuclei enrichment for *sls* mRNAs. Scale bars represent 10 µm. Background stain labels DAPI.

**Figure 4 – figure supplement 1. Nuclear and *sls* mRNA localization**.
(**A**) Molecular Cartography of sparsely expressed genes (*salm* and *CG32121*) at the anterior (right) and middle (left) flight muscle regions labelled in Figure 4A (see also the quantification of nuclear proximity for both genes shown in Figure 3C). Note the proximity of both mRNAs to the nuclei but no mRNA enrichment in anterior regions. (**B**) HCR-FISH imaging of *sls* mRNAs (yellow) in an adult thorax. The dashed line marks the region in which the intensity measurement was achieved. The scale bar represents 50 µm. (C) Quantification of *sls* mRNA localization: intensity profile plots of DAPI (left) and *sls* mRNAs (right) along the anteroposterior axis for 3 sections from 3 different flies indicated in different colors. The positions along the anteroposterior axis were normalized. Note the homogenous DAPI distribution but the terminal *sls* mRNA enrichment. Scale bars represent 10 µm in (A) and 50 µm in (B).

**Figure 5 – figure supplement 1. Variation in mRNA localization using HCR-FISH**.
(A) HCR-FISH of *TpnC4*, A*ct88F*, and *sls* mRNAs in an adult fly thorax. The white boxes mark the zoom-in region shown in (B). (B) Zoom-in of HCR-FISH of *TpnC4*, A*ct88F* (blue in overlay), and *sls* mRNAs in an adult indirect flight muscle. Note that A*ct88F* and *TpnC4* mRNAs largely overlap in this example. The scale bar represents 50 µm in (A) and 10 µm in (B).

**Figure 6 – figure supplement 1: Comparison of head spatial datasets with brain single-cell datasets.**
(**A**) Gene-gene correlation measured across grid-based 5 µm squares and cells. Example mismatches shown in red, matching co-expression in green. (**B**) Composite heatmap showing gene-gene co-expression based on Pearson correlation. Bottom triangle calculated on spatial datasets (using grid-based 5 µm squares). Top triangle calculated using single-cell data. (**C**) Molecular Cartography showing high co-expression of *pros* and *dati*. (**D**) Molecular Cartography showing co-expression of *Indy* and *disco*. (**E**) Molecular Cartography showing expression of *Vmat* and *DAT*. Zoom shows non overlapping expression and expression of *Vmat* outside of nuclei marked by white DAPI. (**F**) Molecular Cartography showing expression of mRNA coding for neuropeptides Ilp2 and Pdf. Zooms show expression outside of nuclei marked by white DAPI signal. (**G**) Stacked bar-plot showing sample composition based on Leiden 1 clustering. Background stain labels DAPI. Scale bars represent 100 µm in (C-F) and 50 µm in the zoomed regions of (E, F).

**Figure 7 – figure supplement 1. Tangram on grid-based quantification**.
(**A**-**D**) Comparison of annotation of spatial data with single-cell RNA-seq. Grid-based squares are colored based on mapped single-cell cluster labels for (A) glia, (B) optic lobe, (C) central brain and (D) uncharacterized clusters using Tangram. In (C) two brain slices are shown at different depths: central (top) and posterior (bottom). (**E**) Effect of manually adjusted

thresholds on Tangram score for nuclei segmentation (left) and grid-based quantification (right) for central brain clusters: 98%-quantile for assigning Pdf-neurons and 99%-quantile for IPC labels. (**F**) Molecular Cartography showing expression of *mirr*. (**G**) Molecular Cartography showing expression of *caup*. Scale bars represent 100 µm. Background stain labels DAPI.

**Table S1**.
All 50 genes used for body MC experiments, including their expression levels and top expressing cell types inferred from the FCA.

**Table S2**.
All 100 genes used for head MC experiments, including their expression levels and top expressing cell clusters inferred from the Pech et al. 2024.

**Table S3**.
Detailed description of the 13 head section samples.

**Table S4**.
All probe sequences used for the HCR-FISH experiments.

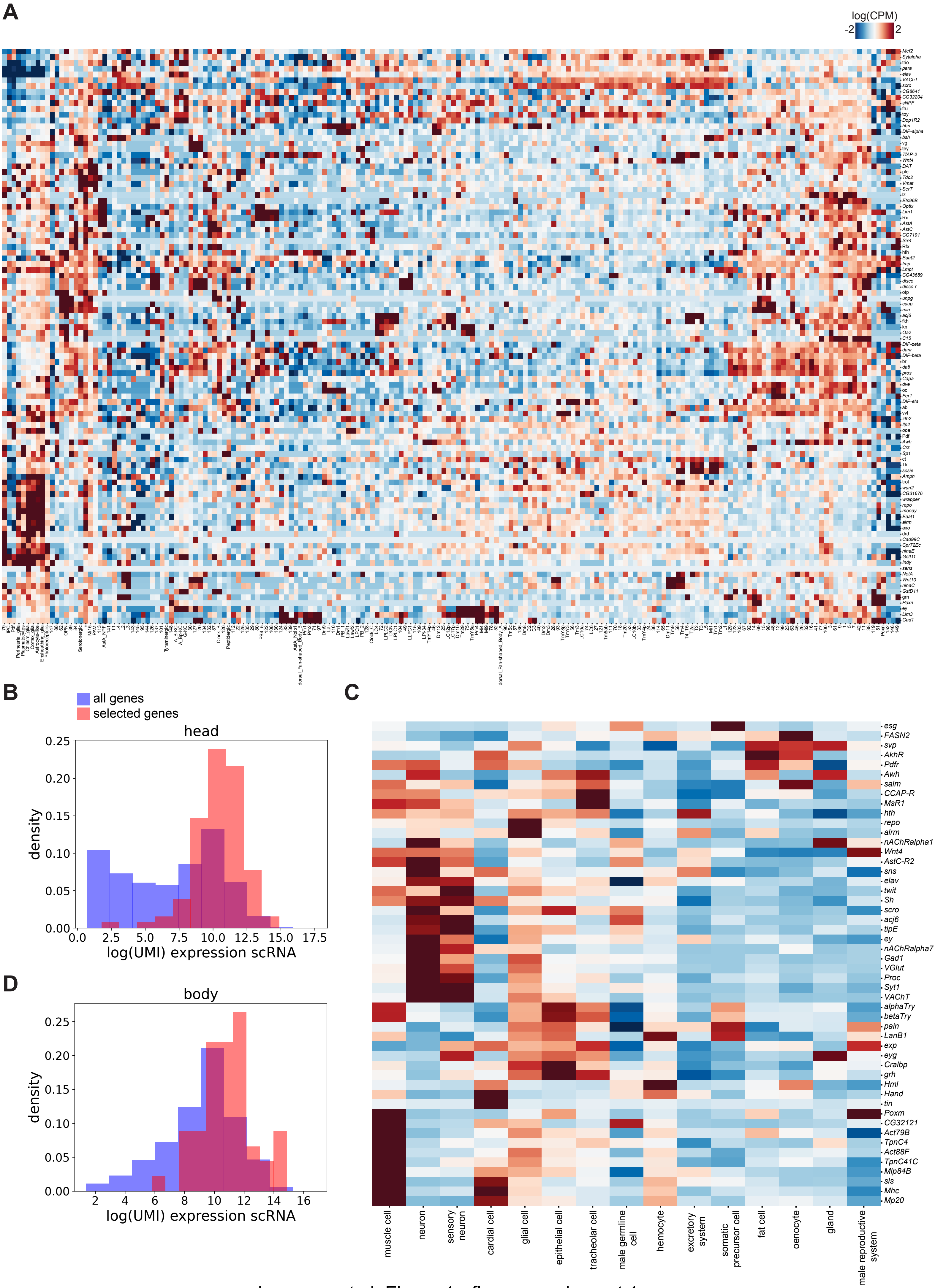


Janssens et al. Figure 1 - figure supplement 1

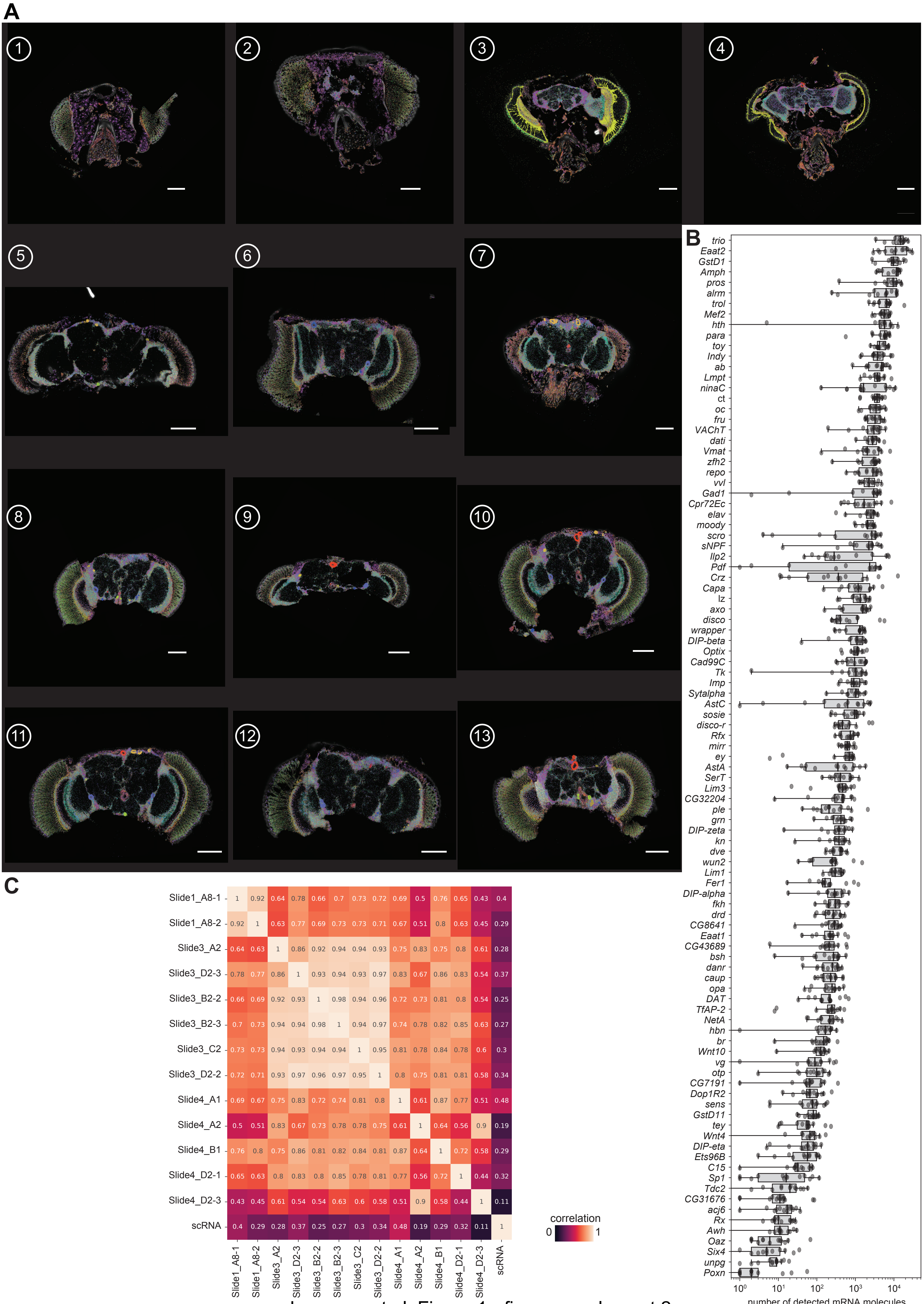


Janssens et al. Figure 1 - figure supplement 2

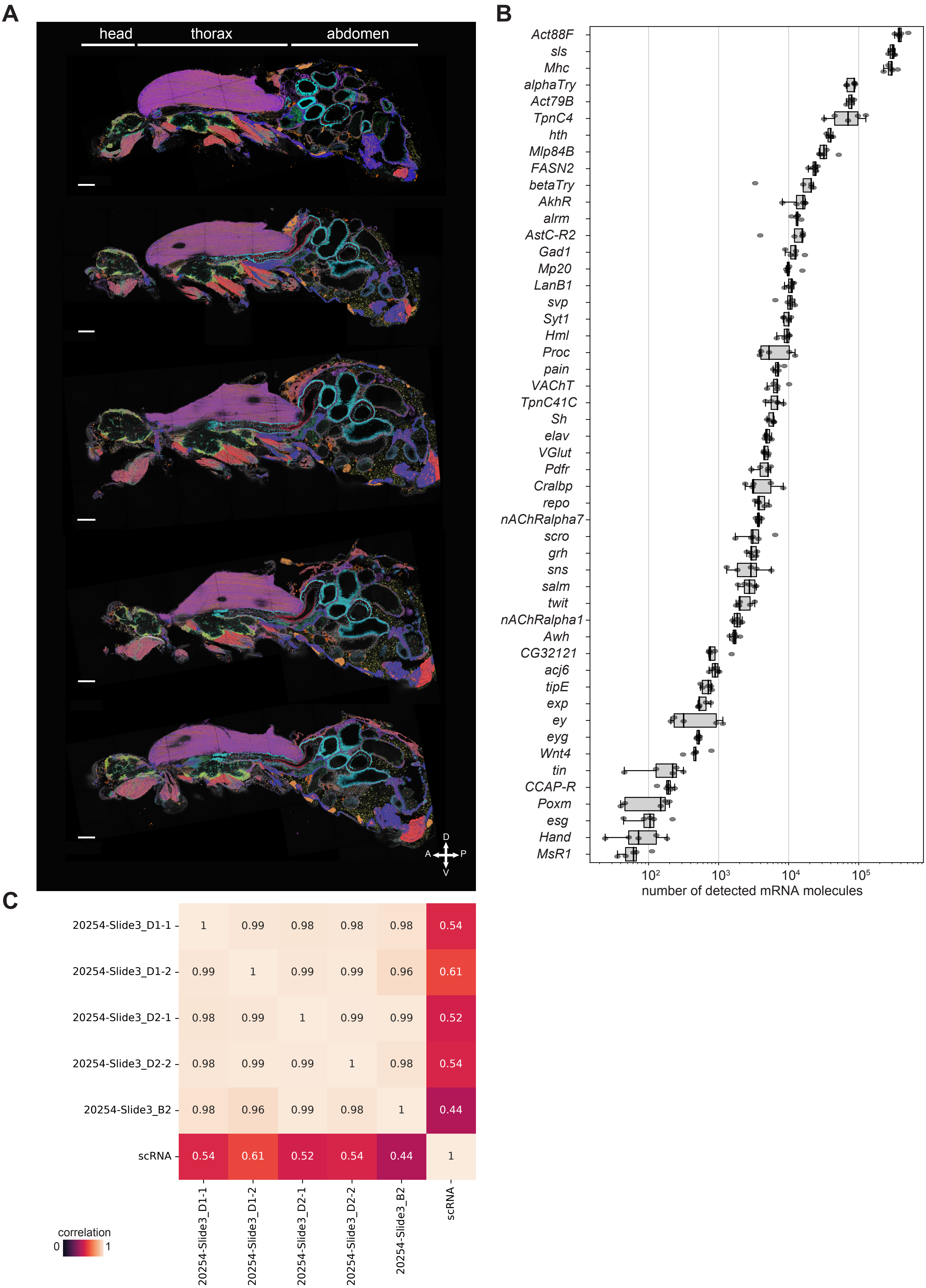


Janssens et al. Figure 1 - figure supplement 3

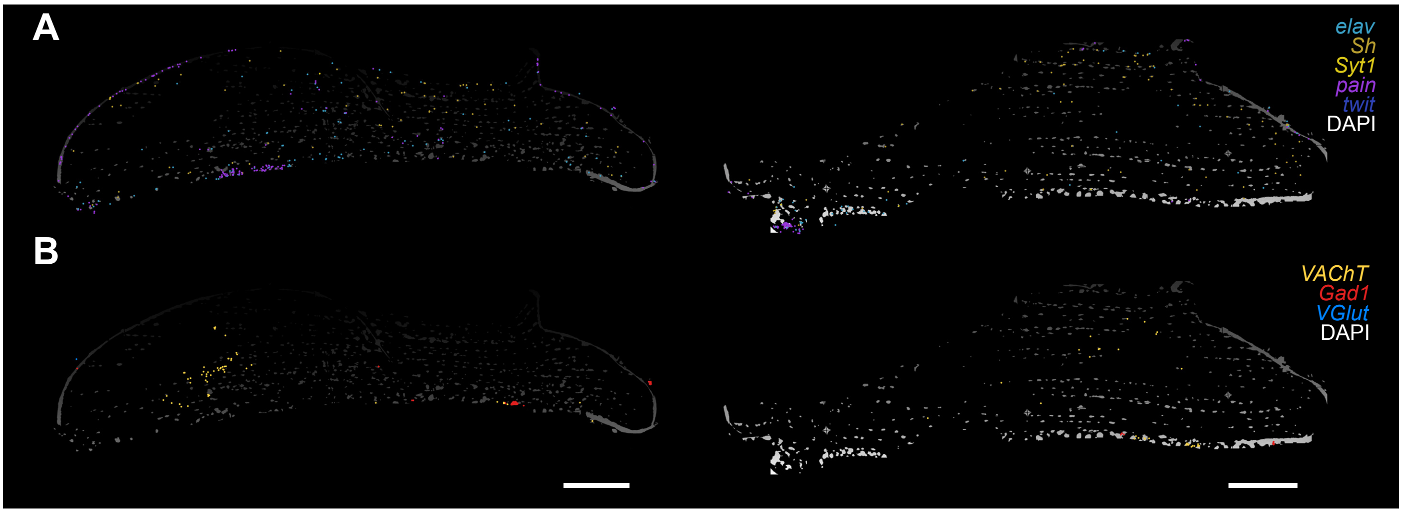


Janssens et al. Figure 2 - figure supplement 1

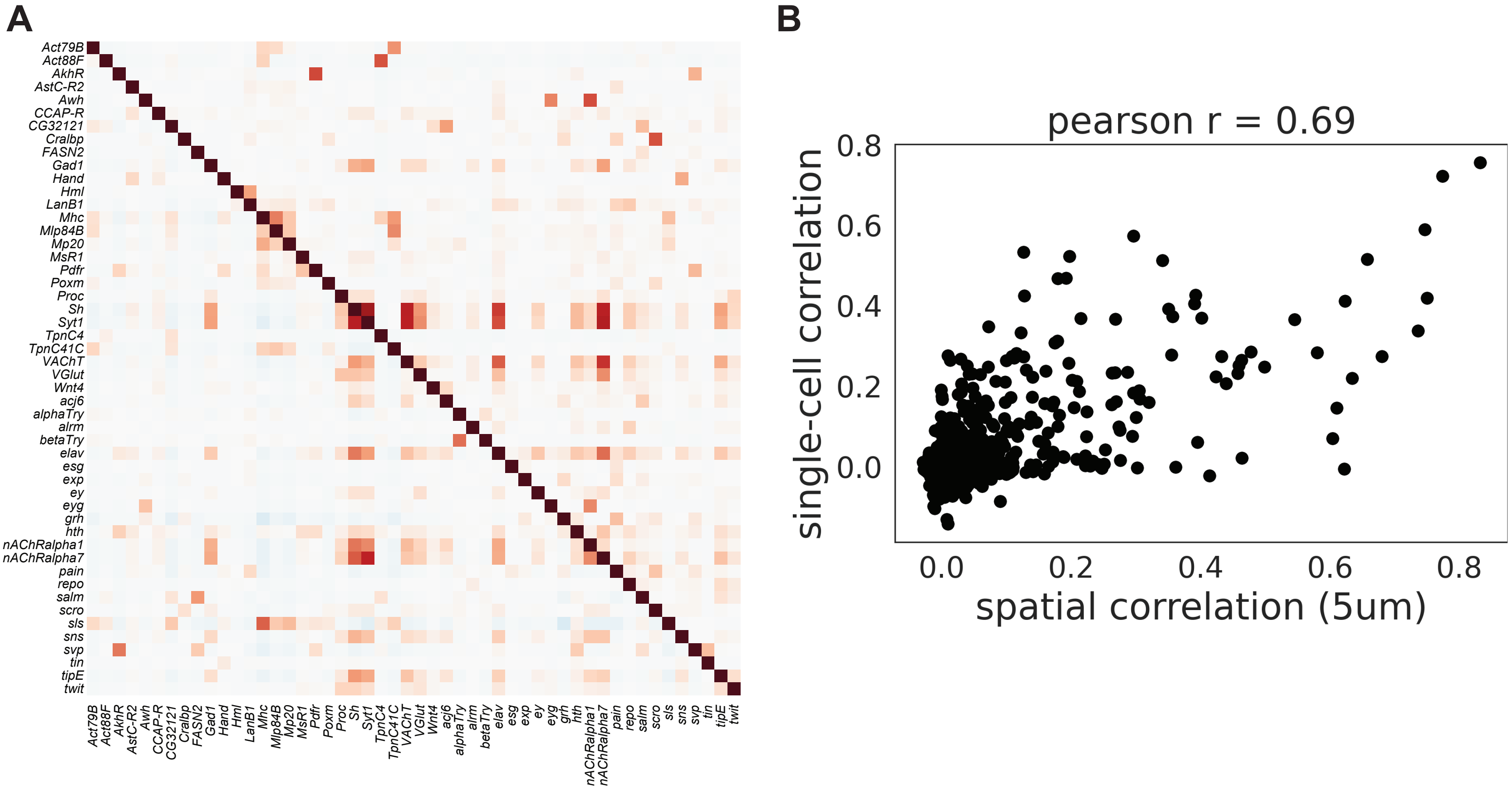


Janssens et al. Figure 2 - figure supplement 2

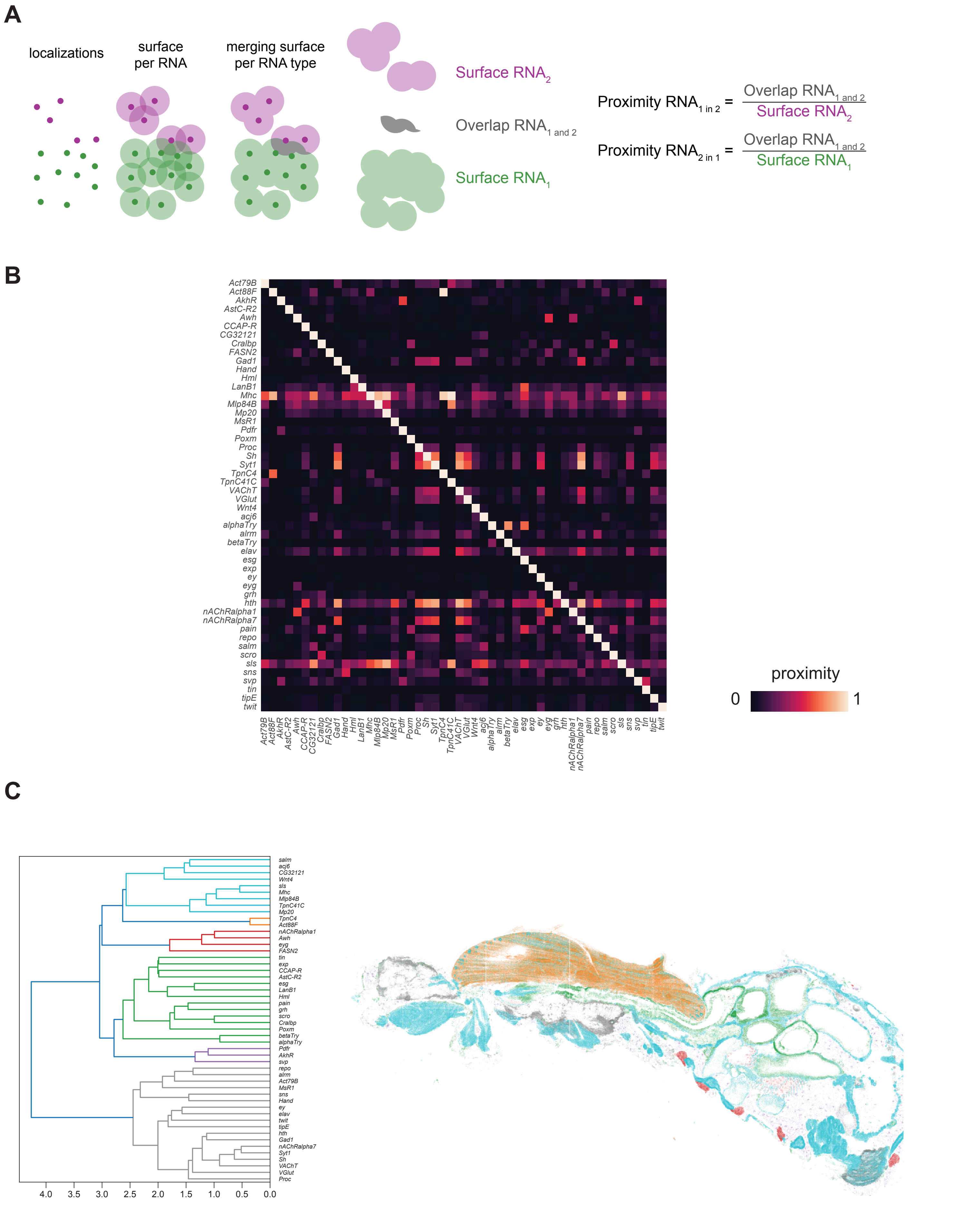


Janssens et al. Figure 2 - figure supplement 3

**A**
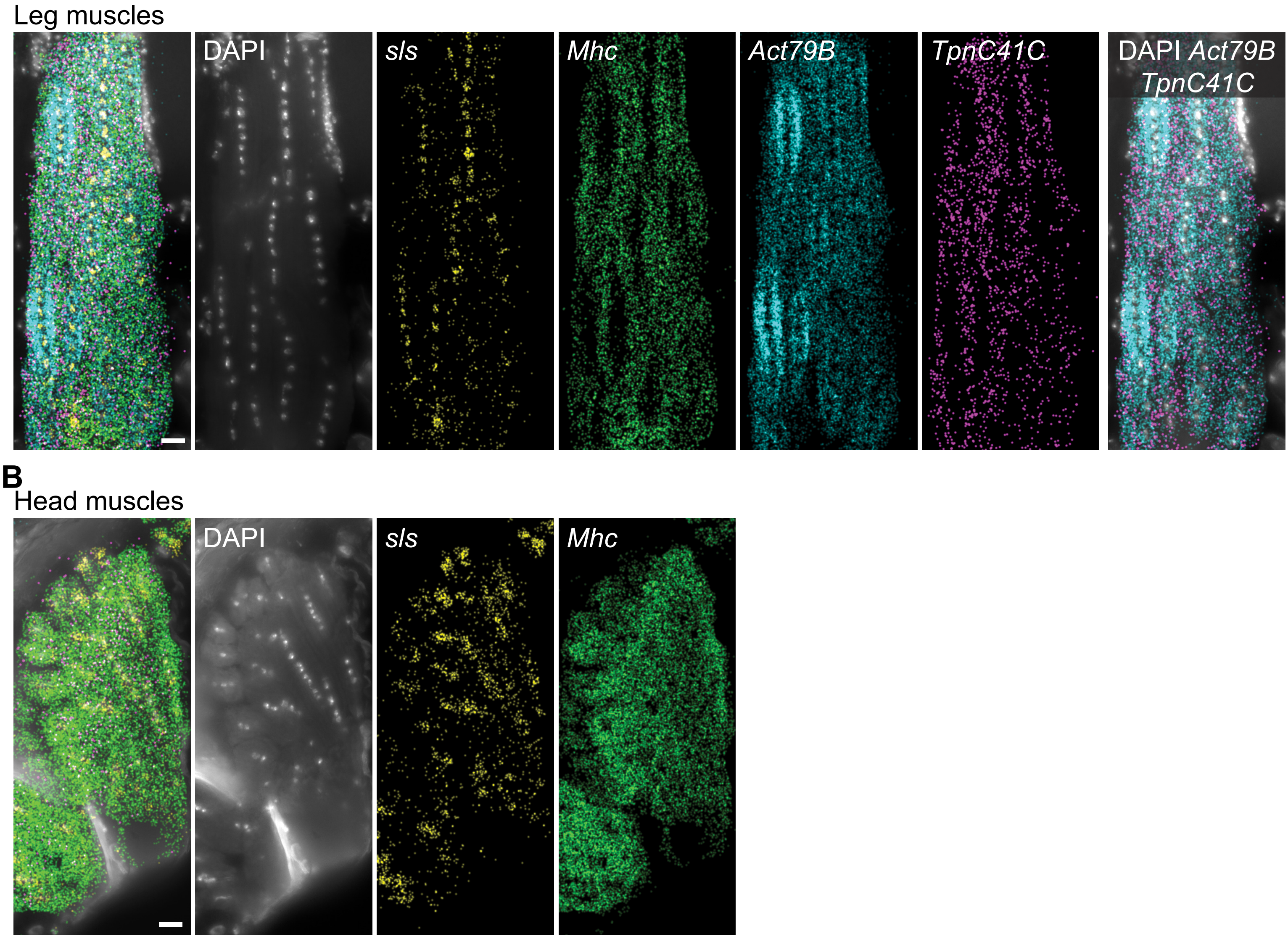


Janssens et al. Figure 3 - figure supplement 1

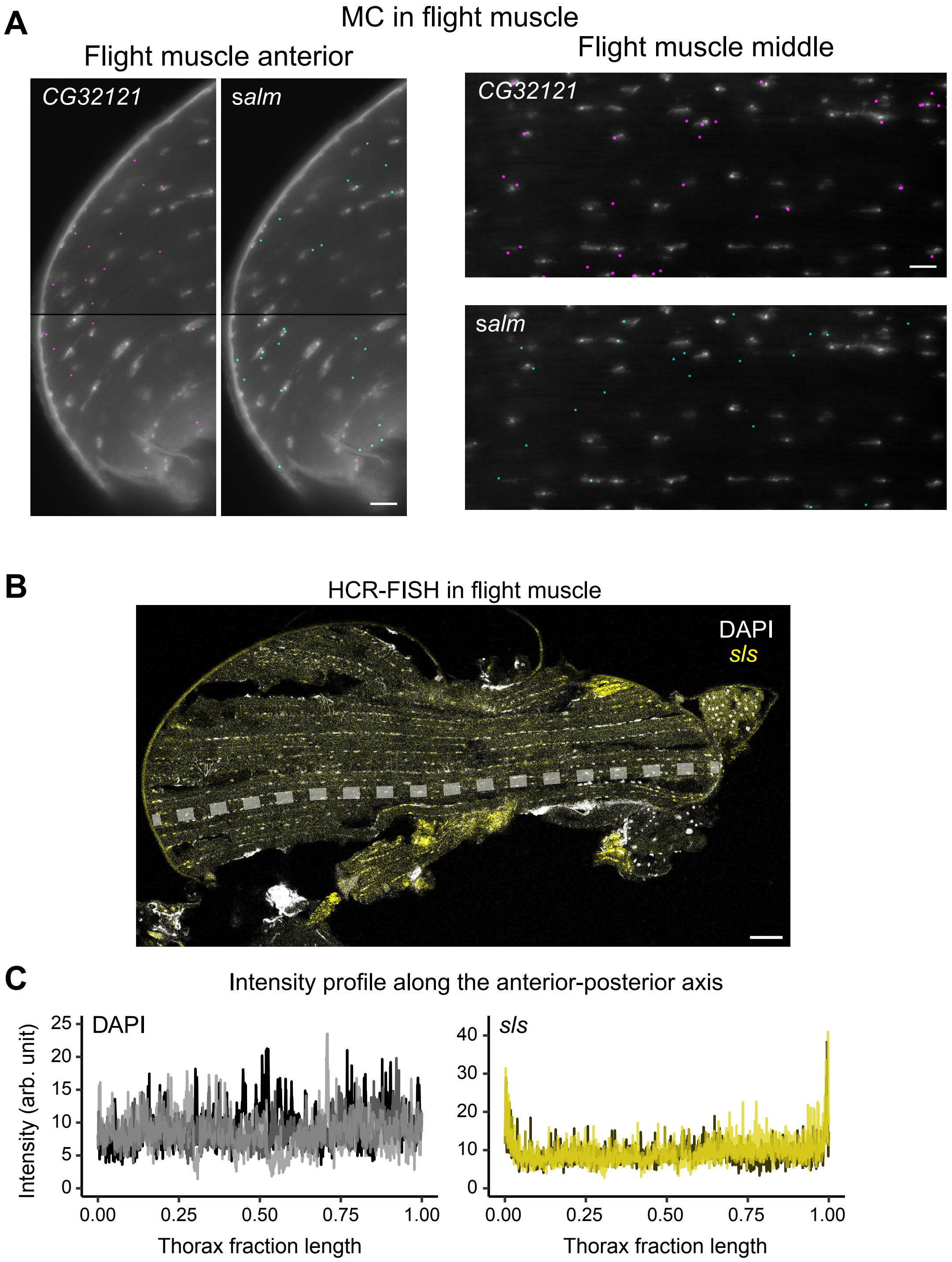


Janssens et al. Figure 4 - figure supplement 1

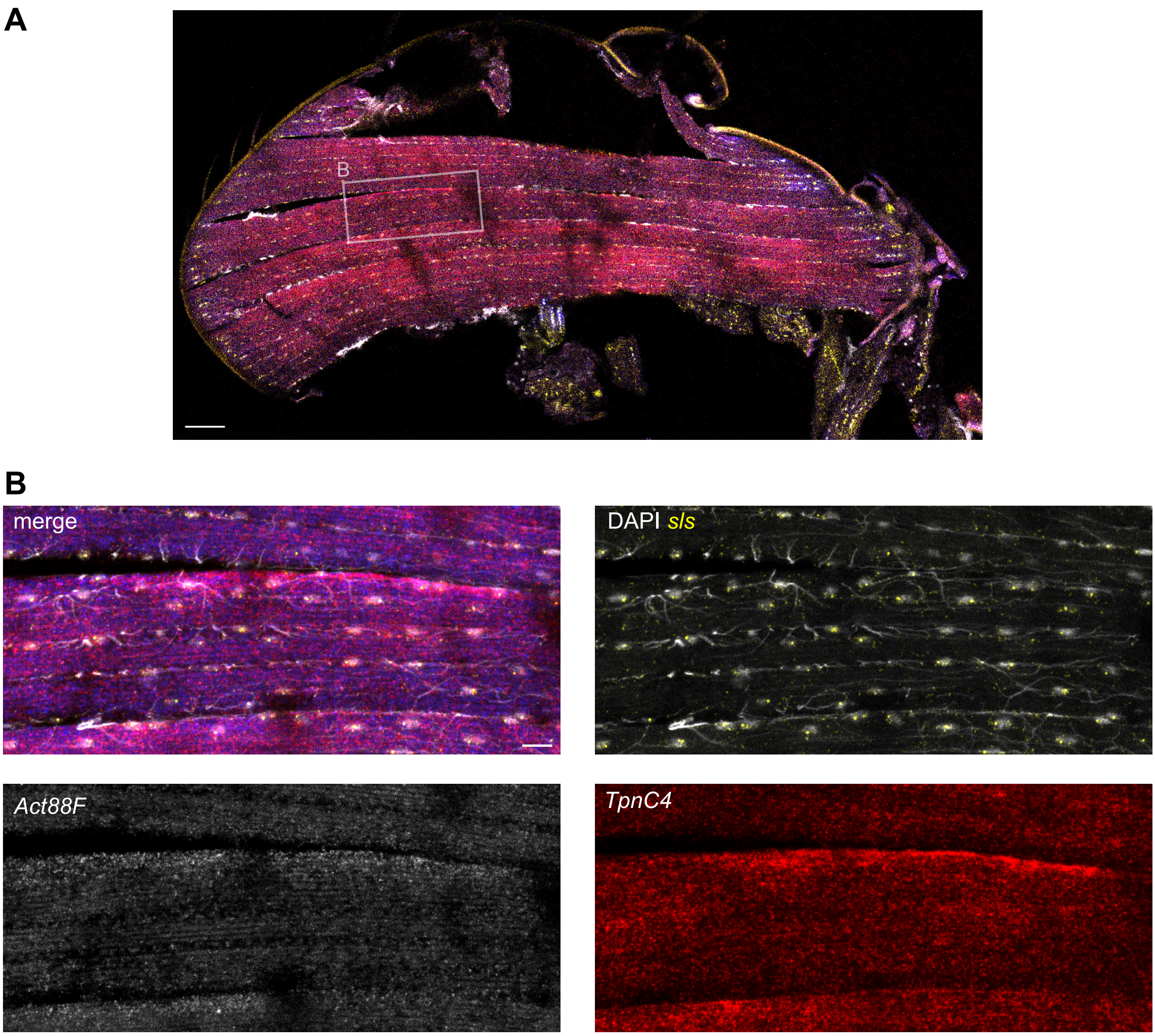


Janssens et al. Figure 5 - figure supplement 1

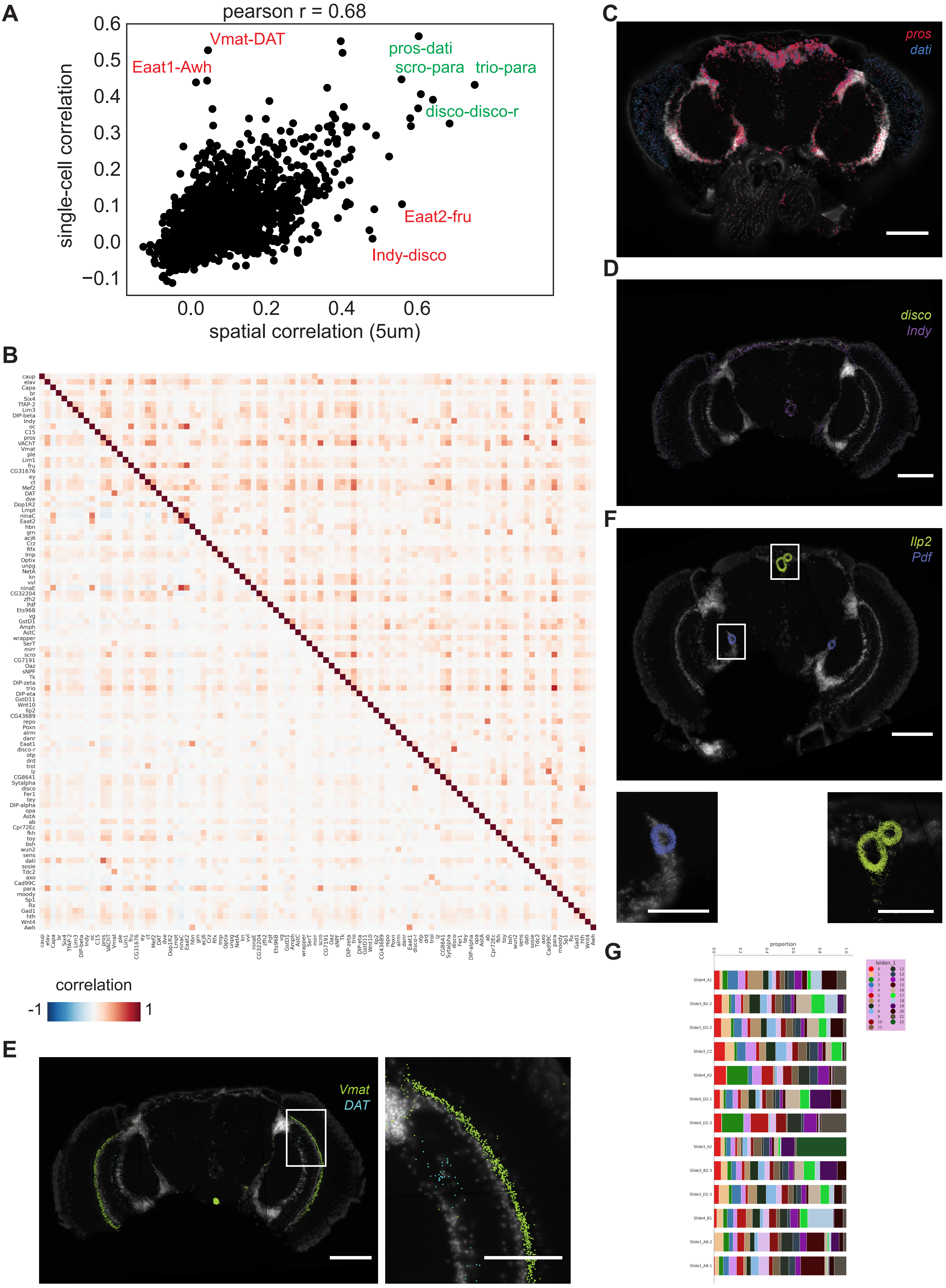


Janssens et al. Figure 6 - figure supplement 1

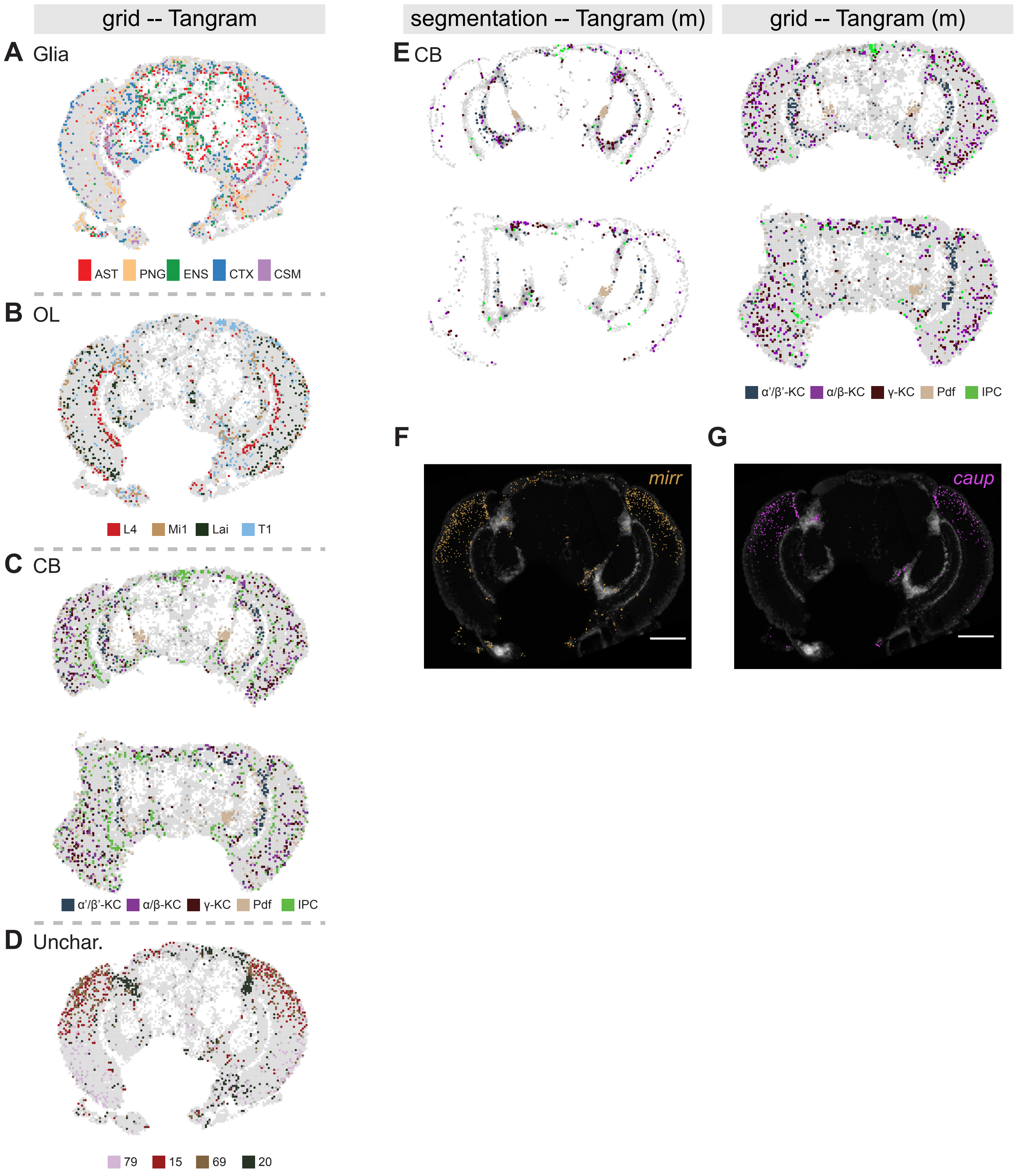


Janssens et al. Figure 7 - figure supplement 1
